# DNA methylation either antagonizes or promotes Polycomb recruitment at transposable elements in Arabidopsis

**DOI:** 10.1101/2024.10.11.617914

**Authors:** Valentin Hure, Maghnia Kawter Elmir, Florence Piron-Prunier, Raphaël Guérois, Angélique Déléris

**Author notes:** Contributed equally (Valentin Hure, Maghnia Kawter Elmir).

## Abstract

Transposable elements (TEs) are usually silenced by DNA methylation and H3K9me2, but in the absence of DNA methylation they can instead acquire the Polycomb-associated mark H3K27me3. Here, we initially set out to test whether DNA methylation and H3K27me3 compete during TE silencing establishment in Arabidopsis. Although we observe this competition at one newly inserted transgenic TE, we remarkably find that H3K27me3 deposition at another neo-inserted TE requires the *de novo* methyltransferase DRM2, revealing that DNA methylation can also promote Polycomb recruitment. Accordingly, genome-wide H3K27me3 profiling in different DNA methylation mutants shows that DNA methylation either antagonizes or promotes H3K27me3 in a locus-specific manner. Targeted DNA methylation experiments as well as the use of a *drm2* catalytic mutant further demonstrate that DNA methylation directly influences Polycomb recruitment. Overall, our work reveals a previously unappreciated interplay between DNA methylation and Polycomb pathways that maintains genome and epigenome integrity in eukaryotes.

## INTRODUCTION

Transposable elements (TEs) are repetitive DNA sequences that have the potential to multiply and move around the genome, thus compromising genome integrity^1^. Nevertheless, this threat is largely mitigated by genome defense mechanisms that efficiently target transposable elements and stably repress their expression across cell division and generations through epigenetic marks such as DNA methylation (5mC). These marks can persist even on TEs that have aged and lost mobility, and such DNA-methylated TEs can play a role as epigenetic regulatory modules that can impact nearby gene regulation^1^. In plants and *Arabidopsis thaliana*, DNA methylation is established and maintained by three classes of DNA methyltransferases. METHYLTRANSFERASE 1 (MET1) maintains CG methylation while CHROMOMETHYLASE 3 (CMT3) and CHROMOMETHYLASE 2 (CMT2) maintain CHG and CHH methylation respectively, and ensure feedback loops with H3K9me2 histone methyltransferases^2^. DOMAINS REARRANGED METHYLASE 2 (DRM2) also maintains non-CG methylation, CHH mainly at TE loci that are smaller than those targeted by CMT2 and CHG to a lesser extent in cooperation with CMT3^3^, and possibly contributes to CG methylation^2,4^. Importantly, DRM2 is able to methylate DNA *de novo* in all sequence contexts as part of the RNA-directed DNA methylation (RdDM) pathway guided by small RNAs (also referred to as small-interfering RNAs or siRNAs)^3^. Thus, *drm2* mutation drastically impairs *de novo* methylation of newly inserted repeats or TE-derived sequences^5,6^, a molecular phenotype also observed to various extents in mutants for RNA interference or RdDM factors connected to DRM2 activity^6,7^.

The detection of a TE that precedes its *de novo* methylation can be achieved in two manners^8^. The first, referred to as “expression-dependent silencing”, is based on TE transcript detection by the RNA interference machinery, which leads to the production of small RNAs^7^. These small RNAs can mediate not only transcript degradation but also RdDM by guiding DRM2 to homologous sequences^3^. Alternatively, if the TE is not transcribed upon insertion but displays sequence homology with another DNA-methylated TE copy that accumulates small RNAs in the genome, then the preexisting small RNAs can similarly guide the *de novo* deposition of DNA methylation by DRM2 by virtue of sequence homology with the TE neo-copies (“homology-dependent silencing”)^8^. The initial DNA methylation levels on a newly introduced repeat-containing transgene or a TE neo-insertion are usually not as high as those of the corresponding endogenous copy, and several generations are needed to reach these levels and a stably repressed state^9,10^.

While very stable across generations and over evolutionary time, the DNA methylation patterns can nonetheless be constitutively pruned by the four members of the DEMETER (DME) family of 5-methylcytosine DNA glycosylases named DME, ROS1 (REPRESSOR OF SILENCING 1), DML2 and DML3 (DEMETER-LIKE 2 and 3). These DNA demethylases act both redundantly and in a locus-specific manner in vegetative tissues to counteract excessive DNA methylation^11–14^. DME (DEMETER), in particular, demethylates transposable elements in companion cells of male and female gametophytes^15,16^ and is required to establish imprints in the endosperm^17^. In contrast to DNA methyltransferases, which have sequence context-dependent activities, DNA glycosylases can demethylate 5mC regardless of the sequence context^12,18,19^. Several functions have been shown for active DNA demethylation, such as preventing the spread of DNA methylation outside of primary targeted sequences^11^ or actively participating in gene control, either at specific stages of development^20^ or by preventing stress-responsive genes from being locked in a constitutively silent state^21,22^.

More recently, TEs have been shown to be decorated by another epigenetic silencing mark, H3K27me3 (trimethylation of lysine 27 of Histone 3)^23^ associated with the evolutionary conserved Polycomb-group (PcG) proteins. This pathway mediates a more dynamic form of transcriptional repression than DNA methylation and has long been thought to specifically target protein-coding genes, particularly development genes for establishing cell identity, or stress-responsive genes^24,25^. H3K27me3 is catalyzed by Polycomb Repressive Complex 2 (PRC2), which is composed of four subunits and is present as different PRC2 cores in Arabidopsis. The catalytic activity is performed by either CURLY LEAF (CLF), SWINGER (SWN) or MEDEA (MEA), which are the three plant homologs of animal ENHANCER OF ZESTE HOMOLOG 2 (EZH2)^23,25^. PRC2 is associated with PRC1, which is composed of two core subunits and mediates the ubiquitination of H2A (H2Aub) to allow a silent yet responsive transcriptional state of H3K27me3-marked targets^26^. In Arabidopsis, the recruitment of PRC2 to genes has been shown to be mediated via Transcription Factors (TFs) that can recognize short recognition sequence motifs called Polycomb responsive elements (PREs), and that were initially described for the recruitment of PRC2 to Drosophila genes^23^. Long noncoding RNAs have also been involved in PRC2 targeting to genes^27,28^. Finally, PRC2 recruitment was shown at some loci to be dependent on the presence of PRC1 and H2Aub^26^.

Interestingly, in recent years, PcG was shown to be the dominant system for silencing TEs in wild-type unicellular eukaryotes and plants from early lineages, thus illustrating the ancient role of PcG in regulating TE silencing^23,29^. However, in Arabidopsis, PcG marks at TEs were shown to vary depending on their context^29^. First, approximately one-third of DNA-methylated TEs can gain H3K27me3 in mutants impaired for DNA methylation, such as *met1* or *ddm1*^30–32^, or in specific cell types that are naturally hypomethylated^33^. This finding implies that DNA methylation, particularly CG methylation, can exclude H3K27me3 deposition at these loci. Second, in some instances, the two marks can co-localize at the molecule level^34–36^, and this was proposed to be constrained by DNA methylation density^34,36^. In this context, the two marks can cooperate in restricting TE activation, for example upon biotic stress^36^. Third, we recently reported that many TE are targeted by the H3K27me3 histone mark in the genome of wild type Arabidopsis plants “instead of” DNA methylation, not only short TE relics but also relatively intact copies^37^. Whether active DNA demethylation can promote H3K27me3 recruitment in some of these cases is unknown.

In this study, we aimed to further investigate the crosstalk between DNA methylation and PcG proteins and reveal novel relationships between these pathways at TEs. Using neo-inserted TE transgenic sequences that are able to recruit both H3K27me3 and DNA methylation *de novo,* we provide evidence that the deposition of H3K27me3 can be either antagonized by DRM2 or, unexpectedly, dependent on DRM2 depending on the neo-inserted TE. This finding sheds light on an unsuspected interconnection between the PcG and RdDM pathways and points to a positive crosstalk between them in that context of TE neo-insertion, in addition to the antagonistic relationship previously proposed. We further identified subsets of endogenous loci that lose H3K27me3 upon impairment of DNA methylation maintenance in *drm2*, but also *met1,* in addition to anticipated subsets where this DNA methylation loss leads to an increase in H3K27me3. These results highlight a dual, locus-specific effect of DNA methylation on PcG recruitment at the genome-wide level which is not restricted to DRM2-dependent DNA methylation. Accordingly, in the *ros1 dml2 dml3* triple mutant, the gain of 5mC leads to the gain of H3K27me3 at some loci but also to the loss of H3K27me3 at others; these findings also reveal a novel function of DNA demethylases in the modulation of H3K27me3 via DNA demethylation. Finally, targeted DNA methylation experiments and the use of *drm2* catalytic mutants demonstrated a direct effect of DNA methylation activity on H3K27me3 deposition. Together, our results uncover new interdependencies between DNA methylation and the PcG machinery, particularly in the context of TE neo-insertion. Thus, the two major silencing pathways, generally thought to be separated, cooperate to shape the epigenome, a concept that could extend to other multicellular eukaryotes.

## RESULTS

### DRM2 is involved in the establishment of H3K27me3 at a newly inserted TE sequence

We previously showed that neo-inserted, transgenic sequences of the mobile *ATCOPIA21* retrotransposon (*AT5TE65370*), consistently recruit H3K27me3 *de novo* in an instructive manner, in a context where position effects are diluted^37^. The endogenous copy is DNA methylated in WT plants and accumulates some 24-nt siRNAs (Supplementary Fig. 1a, left) that are likely to target RNA-directed *de novo* DNA methylation to the homologous transgenic sequences. To verify this, we performed BS-seq on the pools of primary *COPIA21* transformants (TR) used for H3K27me3 analysis^37^. We determined that the transgenic copies are indeed DNA methylated and that they are less DNA methylated than the endogenous copies except in the CHH context (Fig. 1a, Fig. 1b). This higher CHH methylation at transgenes is caused by high *de novo* CHH methylation of the long-terminal repeats (LTR) (Fig. 1b), presumably because small RNAs corresponding to these sequences are particularly abundant in WT plants (Supplementary Fig. 1a, left). As expected, the DNA methylation levels at other TEs in the genome are unchanged upon *ATCOPIA21* sequence insertion (Supplementary Fig. 1b).

**Fig. 1:**
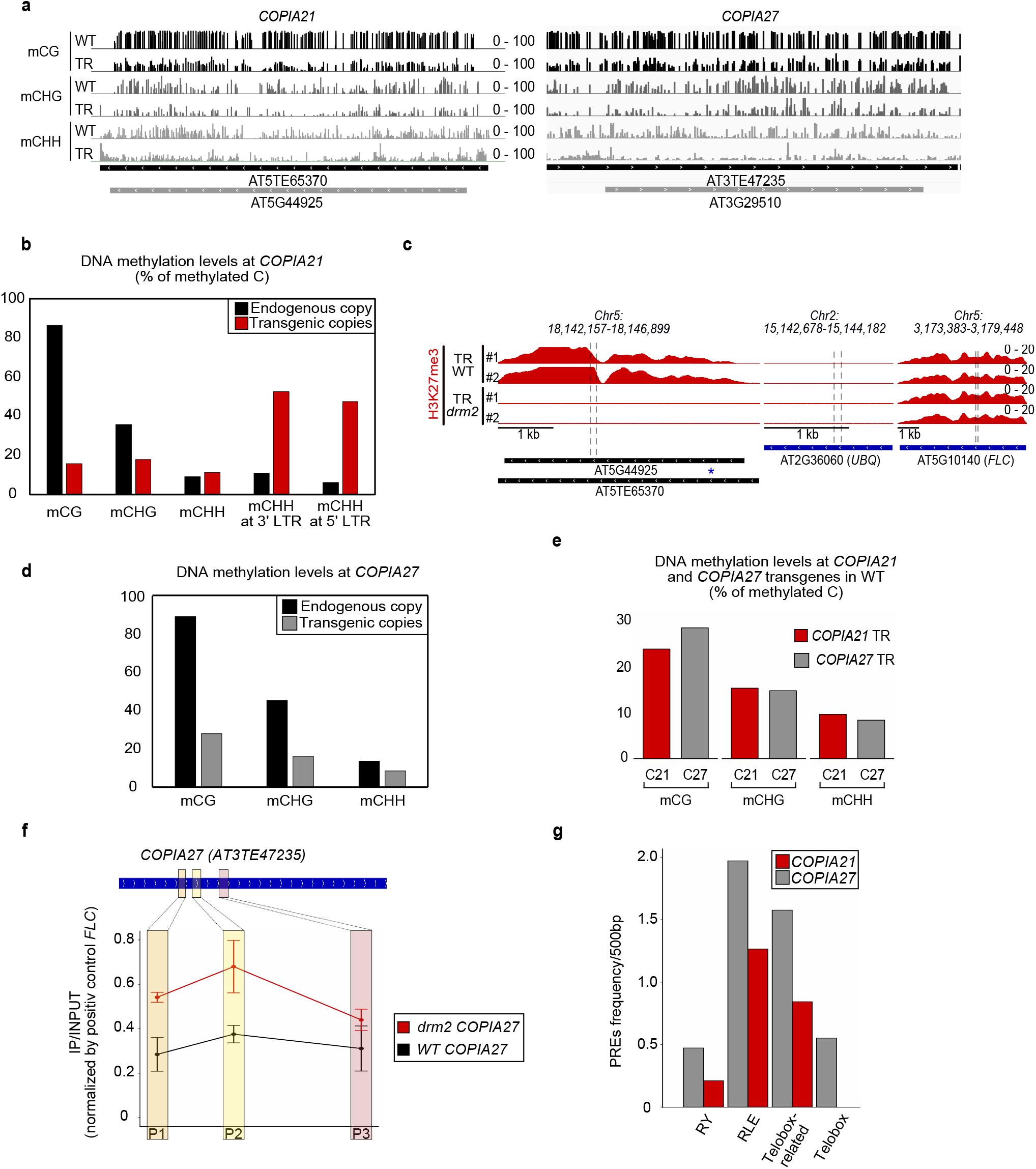
Role of DRM2 in the establishment of *de novo* H3K27me3 at a neo-inserted TE transgenic sequence. **a** Representative genome browser view of DNA methylation status (CG: black, CHG: dark grey, CHH: light grey) at *COPIA21* (left panel) and at *COPIA27* (right panel) in untransformed wild-type (WT) plants and in pools of primary transformants with transgenic TE sequence (TR). **b** Bar plot showing the DNA methylation levels (% of C methylated) in each context at *COPIA21* endogenous copy (black bars) and transgenic copies (red bars). **c** Left panel: Representative genome browser view of H3K27me3 average profile (IP over INPUT) at *COPIA21* sequence in two independent pools of plants transformed with *COPIA21* transgene (#1 and #2) either in WT (Col-0) (upper lanes) or in *drm2* mutant (lower lanes). The H3K27me3 signal in these tracks reflects the H3K27me3 status at the endogene + transgene (SNP as blue star). Middle and right panel: *UBQ* and *FLC* are shown as negative and positive controls respectively for H3K27me3 recruitment. **d** Bar plot showing the DNA methylation levels (% of C methylated) in each context at *COPIA27* endogenous copy (black bars) and transgenic copies (grey bars). **e** Bar plot showing the DNA methylation levels (% of C methylated) in each context at *COPIA21* and *COPIA27* transgenic copies (red and grey bars, respectively) in WT (top). **f** H3K27me3 levels assessed by ChIP-qPCR at *COPIA27* in wild-type and *drm2 COPIA27* transgenic lines. The values from two biological replicates are shown as small horizontal bars, with the mean indicated by a dot. Three different primers pairs were used to address the pattern of H3K27me3 in three different regions of *COPIA27*. No statistical test was performed. **g** Bar plot showing the average number of Polycomb-Responsive Elements (PREs) (per 500bp) at *COPIA21* and *COPIA27* copies (red and grey bars, respectively). Source data are provided as a Source Data file.

To test a model of competition between DNA methylation and H3K27me3 in this context of neo-insertion modelled by transgenic sequences, we transformed the *ATCOPIA21* construct into WT and *drm2* mutant plants impaired in *de novo* DNA methylation. We assessed H3K27me3 levels at the neo-inserted sequences in large pools of primary transformants of the same size for each genetic background. Surprisingly, we observed a drastic and reproducible decrease in H3K27me3 at *ATCOPIA21* copies in the *drm2* mutant plants compared with the WT plants (Fig. 1c, Supplementary Fig. 1c). This finding reveals that DRM2 is required for the *de novo* deposition of H3K27me3 at newly inserted *ATCOPIA21* sequences.

To test if this dependency of H3K27me3 on DRM2-mediated *de novo* DNA methylation is general of all neo inserted TEs, we thought to repeat this experiment with another COPIA element. We chose the *AT3TE47235* copy of the *COPIA27* family which, like *COPIA21(AT5TE65370)* is DNA methylated in WT and marked by H3K27me3 in *ddm1* (Supplementary Fig. 1a). We cloned *AT3TE47235,* transformed WT as well as *drm2* plants and analyzed pools of primary transformants as previously described for pools transformed with *COPIA21*^37^. Consistent with higher levels of small RNAs accumulation at *COPIA27* (*AT3TE47235)* compared to *COPIA21(AT5TE65370)* (Supplementary Fig. 1a, right) and RNA-directed DNA methylation mechanism, *de novo* DNA methylation levels at transgenic *COPIA27* copies were higher than at transgenic *COPIA21*, in particular in the CG context (Fig. 1d and Fig. 1e). H3K27me3 mark was deposited on transgenic *COPIA27* copies in WT plants as expected and observed for other neo-inserted TEs^37^ including *COPIA21* (Fig. 1c, Supplementary Fig. 1c). However, in *drm2*, the levels of H3K27me3 were higher than in WT (Fig. 1f, Supplementary Fig.1d) in contrast with the loss observed at *COPIA21* in *drm2* and in line with our initial competition model. Thus, DRM2 has a dual and differential effect on H3K27me3 deposition on neo-inserted TE sequences. Our BS-seq analyses of the transgenes suggest that this could be linked to the levels of DNA methylation recruited on the neo-inserted sequence. Besides, several PRE-like motifs known to recruit PcG were more abundant in *COPIA27* compared to *COPIA21*, pointing to the possible contribution of sequence composition in the H3K27me3 profiles (Fig. 1g, Supplementary Fig. 1a, red bars).

### DNA methylation is involved in the maintenance of H3K27me3 at endogenous loci

Next, we profiled H3K27me3 in *drm2* mutants and observed that global H3K27me3 levels at genes and endogenous TEs were essentially unchanged (Supplementary Fig. 2a). Thus, DRM2 is not required for overall PRC2 activity and does not appear to indirectly affect Polycomb function. However, closer data inspection revealed two contrasting behaviors at specific loci. A small subset of loci showed increased H3K27me3 in *drm2* (Fig. 2a,b; Supplementary Fig. 2b). At these loci, DNA methylation normally prevents H3K27me3 deposition. We therefore refer to them as **LOCK** (*Loci of Opposition between Cytosine methylation and K27me3*). This supports the idea that DRM2-dependent DNA methylation can antagonize H3K27me3, consistent with effects previously described for CG methylation^31,32^. Of note, the extent of H3K27me3 gain did not seem proportional to the extent of CHH loss, possibly because the loss of a few critical cytosines could be sufficient to allow H3K27me3 deposition (Supplementary Fig. 2b, yellow regions and arrows).

**Fig. 2:**
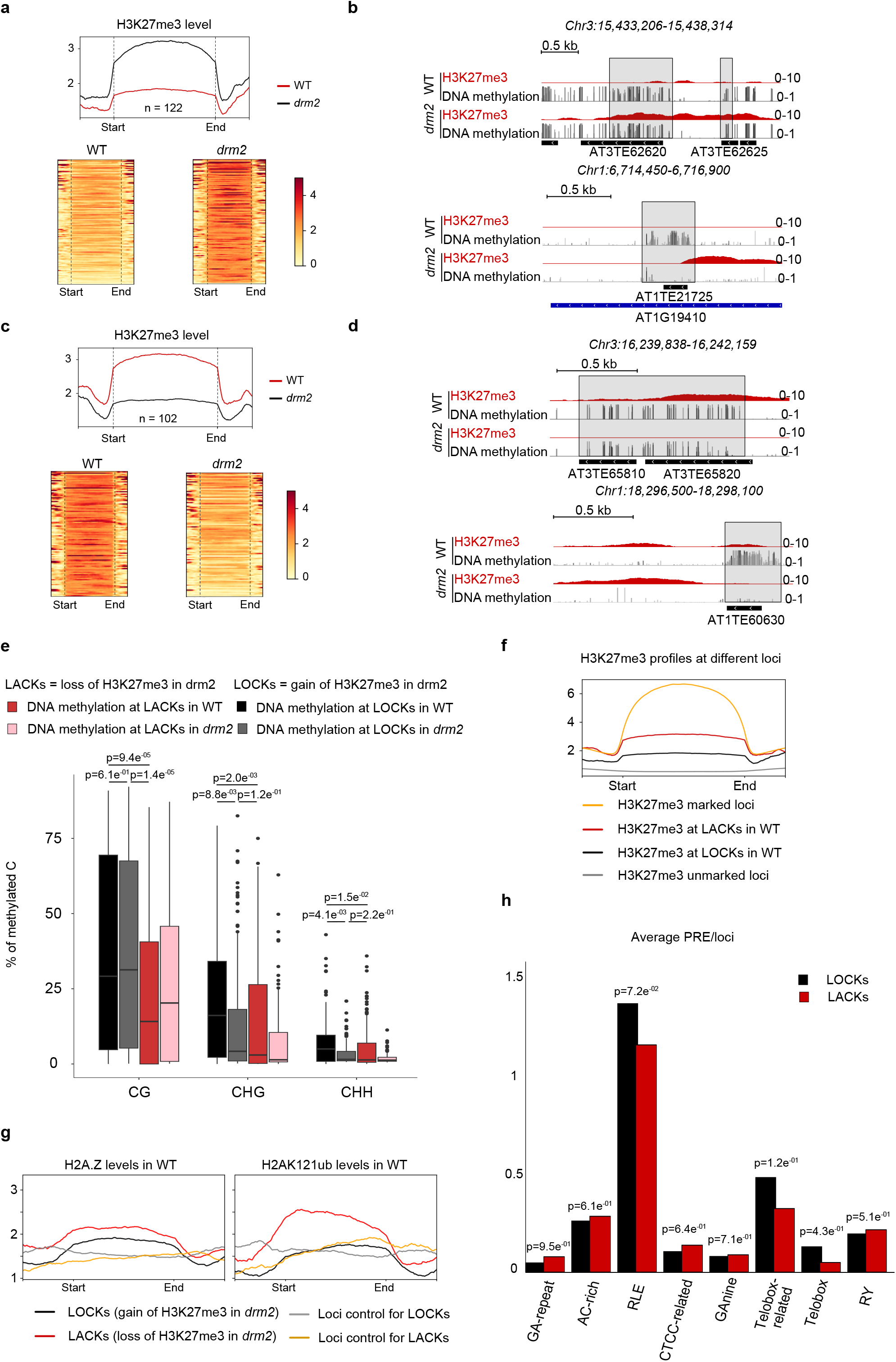
Role of DNA methylation in the recruitment of H3K27me3 at endogenous loci. **a** Heatmaps and metagenes showing H3K27me3 levels for loci that gain H3K27me3 peaks in *drm2* compared to WT (LOCKs). **b** Representative genome browser views showing H3K27me3 ChIP-seq and DNA methylation BS-seq data in WT and *drm2*. Published differentially methylated regions (DMRs) are shown with grey boxes (*Stroud et al, 2013*). **c** Heatmaps and metagenes showing H3K27me3 levels for loci that loose H3K27me3 peaks in *drm2* compared to WT (LACKs). **d** Representative genome browser views showing H3K27me3 and DNA methylation data in WT and *drm2*. Loss of H3K27me3 in *drm2* compared to WT is associated with previously published DMRs (grey boxes) (*Stroud et al, 2013*). **e** Boxplot showing DNA methylation in three contexts for different H3K27me3 peak regions (i.e., peak+500bp on the sides to account for any effect of DNA methylation patches on peak edges). Black and grey bars show the DNA methylation levels in the same regions, in WT and *drm2*, respectively. These regions correspond to peaks of H3K27me3 gained in *drm2* (n=122). Red and pink bars show the DNA methylation levels in the same regions, in WT and *drm2*, respectively. These regions correspond to peaks of H3K27me3 lost in *drm2* (n=102). Each data point represents a genomic locus. Boxplots show the median, with boxes indicating the 25th and 75th percentiles and whiskers extending to values within 1.5× the interquartile range. **f** Metaplots showing H3K27me3 profiles at loci marked by H3K27me3 (orange), loci loosing H3K27me3 in *drm2* (LACKs) (red), loci marked by H3K27me3 gained in *drm2* (LOCKs) (black) and loci not marked by H3K27me3 are shown as a negative control (grey). **g** Metaplots showing H2A.Z variant and H2AK121ub profiles at LOCKs (loci that gain H3K27me3 in *drm2*, black), at LACKs (loci that loose H3K27me3 in *drm2*, red) and at associated control regions (random loci of the same length). **h** Bar plot showing the average number of Polycomb-Responsive Elements (PREs) per locus (normalized by the size of the locus). LOCKs and LACKs are shown in black and red, respectively. Statistical analyses were performed using a two-sided Wilcoxon-Mann-Whitney U-test, p-values are indicated. Source data are provided as a Source Data file.

On the other hand, another subset of loci showed a strong reduction or complete loss of H3K27me3 in *drm2* (Fig. 2c,d; Supplementary Fig. 2c), similar to our observations at the *ATCOPIA21* transgene. At these loci, DRM2-dependent DNA methylation normally *promotes* H3K27me3. We therefore refer to them as **LACK** (*Loci of Association between Cytosine methylation and K27me3*). Their loss of non-CG methylation in *drm2* confirms that they are direct DRM2 targets (Fig. 2e). Again, the degree of H3K27me3 loss was not proportional to CHH loss, suggesting that reduced methylation at key cytosines is sufficient to destabilize PRC2 occupancy (Supplementary Fig. 2c, yellow regions and arrows). Together, these observations show that DRM2-mediated non-CG methylation is required at a subset of loci to maintain proper H3K27me3 patterns.

To understand why H3K27me3 responds differently at these two classes of loci, we compared their DNA methylation, chromatin features, and other intrinsic properties. LACK (where H3K27me3 marks are promoted by DRM2 and co-localize with DNA methylation) (Fig. 2e, red bars), are significantly less DNA methylated, in all three sequence contexts than LOCK (where H3K27me3 marks are antagonized by DRM2 )(Fig. 2e, black bars). This suggests that a certain threshold of DNA methylation may contribute to determine whether H3K27me3 deposition is antagonized (high DNA methylation) or promoted (low DNA methylation) by DRM2. In support of this, non-CG methylation levels at loci that gain H3K27me3 in *drm2* (LOCK) match wild-type levels at LACK (Fig. 2e, grey bars versus red bars). Besides, H3K27me3 enrichment at LACK is lower than at typical H3K27me3 regions but higher than at LOCK (Fig. 2f). Additional meta analyses using published datasets^38–40^ showed that LACK are generally depleted in small RNAs and constitutive heterochromatin marks (H2A.W, H3K9me1 and H3K9me2) compared to LOCK (Supplementary Fig. 2d,e). Instead, LACK are rather enriched in hallmarks of facultative heterochromatin marks and of responsive genes including H2A.Z histone variant, H2Aub and H3/H4ac chromatin marks (though to a low level for the latest) compared to LOCK (Fig. 2g, Supplementary Fig. 2e,f). Besides, LACK tend to be shorter than LOCK (Supplementary Fig. 2g), suggesting they may represent older degenerated TEs and consistent with shorter TEs exhibiting reduced DNA methylation ^41^. None of these classes show a bias toward specific chromosomal regions (Supplementary Fig. 2h) nor a strong enrichment for specific TE superfamilies (Supplementary Fig. 2i). Finally, consistent with transgene analyses, LACK contain significantly fewer Polycomb response elements (PREs) than LOCK (Fig.2h). Together, these analyses point to particular epigenetic and genetic signatures of LACK versus LOCK which likely contribute to explain the locus-specific and opposite effects of DRM2 on H3K27me3 deposition.

Finally, to determine whether PRC2 recruitment depends specifically on DRM2 or whether other DNA methyltransferases can serve similar roles, we profiled H3K27me3 in *met1* mutant plants. As previously reported^31^, loss of MET1 leads to H3K27me3 gain at many MET1 targets. In our framework, these correspond to MET1-LOCK, loci where MET1-dependent CG methylation normally prevents H3K27me3. Some of these MET1-LOCK overlap with DRM2-LOCK, while others are MET1-specific (Supplementary Fig. 3a,b), indicating that both MET1 and DRM2 can generate regions where DNA methylation excludes PRC2. On the other hand, we also identified MET1-targets that lose H3K27me3 in *met1*, corresponding to MET1**-**LACK, where MET1-dependent DNA methylation promotes H3K27me3. These are largely distinct from the DRM2-LACK where H3K27me3 levels at DRM2-LACK remain unchanged in *met1* (Supplementary Fig. 3c,d). Thus, DRM2 and MET1 act on largely non-overlapping sets of loci to promote H3K27me3. Finally, DRM2-LACK and MET1-LACK show minimal overlap with TEs upregulated in *drm2 and met1*—less than 2% and 10% respectively: this shows that loss of H3K27me3 at LACK is largely not a consequence of transcriptional reactivation (Supplementary Fig. 3e). Together, these results show that PRC2 recruitment and activity can be influenced either by DRM2 or by other DNA methyltransferases, depending on the locus.

### DNA demethylation impacts H3K27me3 patterning at endogenous loci

To further explore the interconnections between DNA methylation and PRC2 activity, we profiled H3K27me3 in the triple DNA demethylase mutant *rdd* (*ros1 dml2 dml3).* At a subset of loci normally marked by H3K27me3 in WT and targeted by ROS1/DML2/DML3^42^, increased DNA methylation in *rdd* resulted in a loss of H3K27me3 (Fig. a,b and Supplementary Fig. 4a). This finding indicates that H3K27me3 recruitment at some TEs could be promoted by active demethylation. We refer to these loci as RDD-LOCK, where DNA hypermethylation antagonizes PRC2, consistent with the previously reported inhibitory effect of DNA methylation on H3K27me3. Interestingly, and reminiscent of the DRM2-dependent behavior, we also identified loci, that are weakly or not marked by H3K27me3 in WT and gain H3K27me3 upon DNA hypermethylation in *rdd* (Fig. 3c,d and Supplementary Figure 4b). We refer to these loci as RDD-LACK, where DNA hypermethylation promotes PRC2 recruitment.

**Fig. 3:**
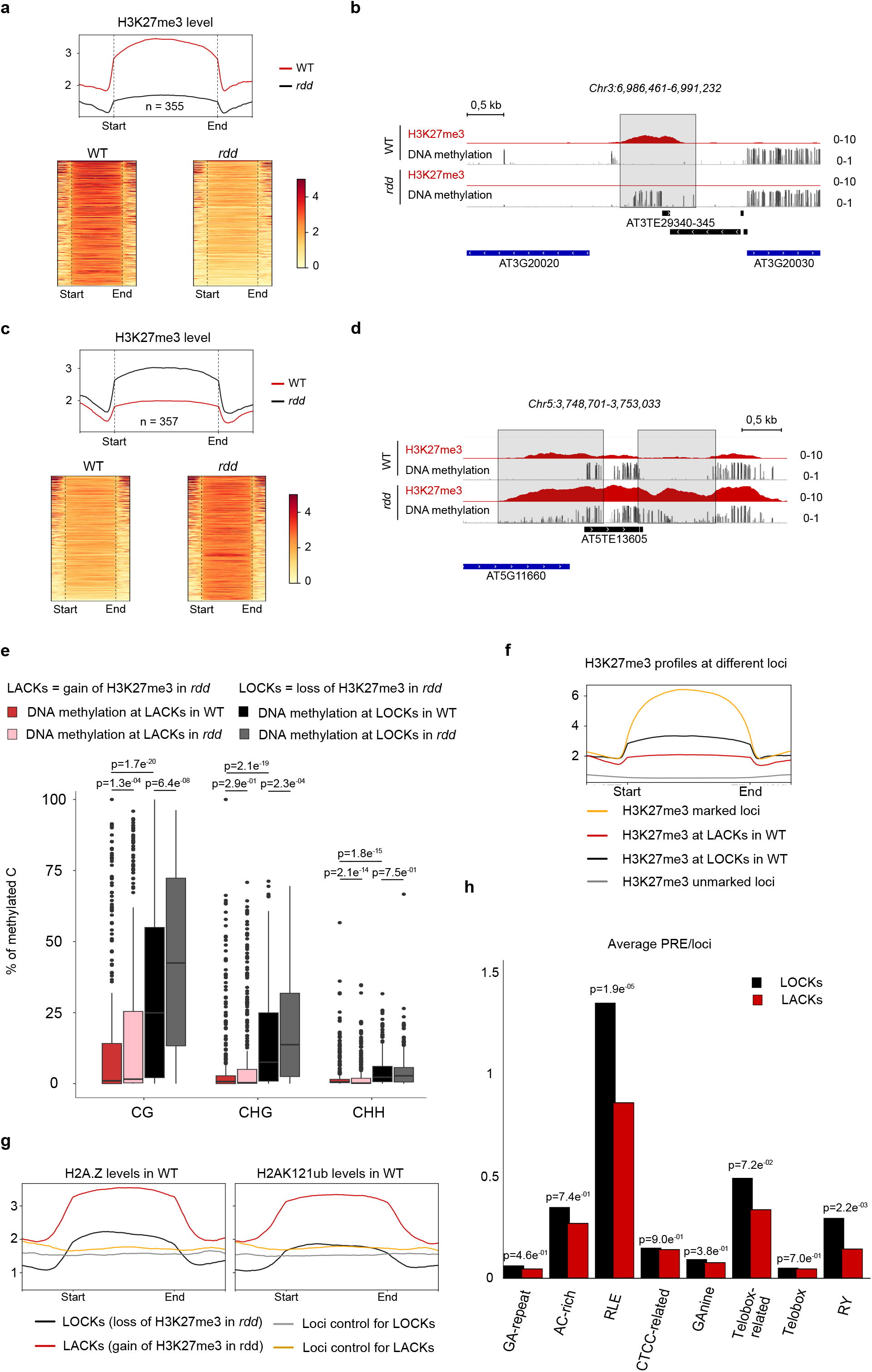
DNA demethylation impacts H3K27me3 patterning at endogenous loci.a. Heatmaps and metagenes showing H3K27me3 levels for loci that loose H3K27me3 peaks in *rdd* compared to WT (LOCKs). **b** Representative genome browser views showing H3K27me3 ChIP-seq and DNA methylation BS-seq data in WT and *rdd*. Published differentially methylated regions (DMRs) are shown with grey boxes (*Duan et al, 2015*). **c** Heatmaps and metagenes showing H3K27me3 levels for loci that gain H3K27me3 peaks in *rdd* compared to WT (LACKs). **d** Representative genome browser views showing H3K27me3 and DNA methylation data in WT and *rdd*. Gain of H3K27me3 in *rdd* compared to WT is associated with previously published DMRs (grey boxes) (*Duan et al, 2015*). **e** Boxplot showing DNA methylation in three contexts for different H3K27me3 peak regions (i.e., peak+500bp on the sides to account for any effect of DNA methylation patches on peak edges). Black and grey bars show the DNA methylation levels in the same regions, in WT and *rdd*, respectively. These regions correspond to peaks of H3K27me3 lost in *rdd* (n=355). Red and pink bars show the DNA methylation levels in the same regions, in WT or in *rdd*, respectively. These regions correspond to peaks of H3K27me3 gained in *rdd* (n=357). Each data point represents a genomic locus. Boxplot show the median, with boxes indicating the 25th and 75th percentiles and whiskers extending to values within 1.5× the interquartile range. **f** Metaplots showing H3K27me3 profiles at loci marked by H3K27me3 (orange), loci loosing H3K27me3 in *rdd* (LOCKs) (black), loci marked by H3K27me3 gained in *rdd* (LACKs) (red) ; loci not marked by H3K27me3 are shown as a negative control (grey). **g** Metaplots showing H2A.Z variant and H2AK121ub profiles at LOCKs (loci that loose H3K27me3 in *rdd*, black), at LACKs (loci that gain H3K27me3 in *rdd*, red) and at associated control regions (random loci of the same length). **h** Bar plot showing the average number of Polycomb-Responsive Elements (PREs) per locus (normalized by the size of the locus). LOCKs and LACKs are shown in black and red, respectively. Statistical analyses were performed using a two-sided Wilcoxon-Mann-Whitney U-test, p-values are indicated. Source data are provided as a Source Data file.

As above, to understand why H3K27me3 responds differently at these two classes of loci, we compared their DNA methylation, chromatin features, and other intrinsic properties. Similar to the DRM2-defined classes, RDD-LACK exhibit significantly lower methylation in all sequence contexts than RDD-LOCK (Fig. 3e) and show reduced small RNA abundance (Supplementary Fig. 4c). These observations again suggest that a DNA methylation threshold helps determine whether PRC2 deposition is antagonized (higher methylation at LOCK) or promoted (lower methylation at LACK). Notably, RDD-LOCK exhibit intermediate levels of both DNA methylation and H3K27me3 (Fig. 3e,f), placing them between fully marked and unmarked chromatin states.

To further characterize these subsets, we performed meta-analyses to examine additional chromatin and genomic features of these subsets. RDD-LOCK are enriched in constitutive heterochromatin marks (H2A.W, H3K9me1, H3K9me2) (Supplementary Fig. 4d), whereas RDD-LACK are enriched in hallmarks of facultative or dynamic chromatin, including H2Aub, H2A.Z, and H3/4 acetylation marks (Fig. 3g; Supplementary Fig. 4e), consistent with known ROS1-associated chromatin environments^43,44^. At the sequence level, RDD-LOCK contain more PRE motifs (RY, RLE, Telobox) (Fig. 3h), whereas RDD-LACK are shorter and exhibit higher GC content and elevated frequencies of CG, CHG, and CHH motifs (Supplementary Fig. 4f). Both classes are distributed across pericentromeric and arm regions and show no strong bias toward specific TE superfamilies (Supplementary Fig. 4g,h).

Importantly, RDD-LACK and RDD-LOCK are distinct from the corresponding DRM2-defined classes, consistent with their fundamentally different behaviors in wild type: DRM2-LACK are H3K27me3-marked and lose H3K27me3 in *drm2*, whereas RDD-LACK are H3K27me3-unmarked (or weakly marked) and gain H3K27me3 in *rdd*; conversely, DRM2-LOCK are H3K27me3-unmarked (or weakly marked) and gain H3K27me3 in *drm2*, whereas RDD-LOCK are H3K27me3-marked and lose it in *rdd*.

Altogether, these analyses reveal that active DNA demethylation can either promote or antagonize PRC2 recruitment, generating LACK and LOCK behaviors analogous to those observed with the DNA methylation pathway. The opposing outcomes at these loci are associated with distinct genetic and epigenetic signatures shared across the corresponding DRM2- and RDD-defined classes, and point to locus-specific rules that govern the interplay between DNA methylation dynamics and Polycomb targeting.

### A targeted DNA methylation approach shows the direct impact of DNA methylation on H3K27me3 deposition in a locus-dependent manner

We next wanted to verify whether the loss of H3K27me3 at the DNA hypermethylated regions was the direct consequence of ectopic DNA methylation. For this purpose, we constructed an inverted repeat transgene to produce small RNAs able to override the dominance of ROS1 activity over RdDM^22^ and induce DNA methylation^45^ at a specific TE locus (*AT1TE59770*, i.e LOCK Target 1) (Fig. 4a left panel, Fig. 4b). In the RNAi lines that we isolated, we observed a gain of DNA methylation in all three sequences contexts (Fig. 4c left panel) and this led to a clear decrease in H3K27me3 at the endogenous TE (“Target 1”) compared to the transgenic lines with an unrelated transgene (Fig. 4d and Supplementary Fig. 5b, left panels). We thus show for the first time that targeting small RNA-directed DNA methylation to a H3K27me3-marked locus can cause a direct loss of H3K27me3.

**Fig. 4:**
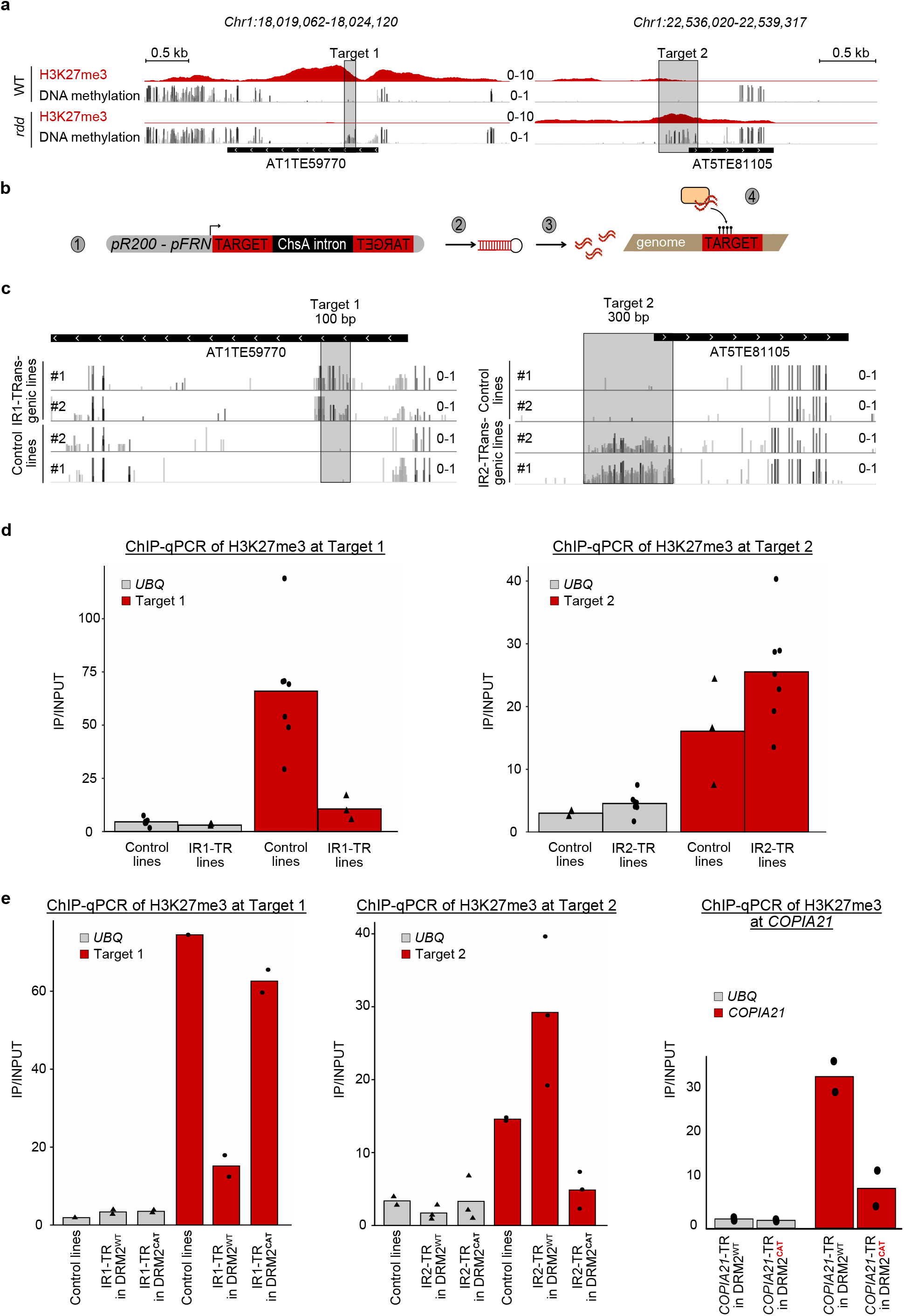
Targeted DNA methylation by Inverted-Repeats (IR) reveals a dual role of DNA methylation on H3K27me3 patterning at endogenous TEs. a Representative genome browser view showing H3K27me3 and DNA methylation in WT and *rdd* mutant. The regions highlighted in grey correspond to DMRs that were cloned for the experiments shown in panels 4C and 4D. b Experimental design: (1) Two TE target regions were selected (Target 1 and 2); for each target, the sequence was cloned as an Inverted Repeat (IR) separated by the ChsA intron in the pR200-pFRN vector. (2) Expression of the IR transgene leads to hairpin formation. (3) Hairpins are processed by DICER-like proteins into small RNAs. (4) Small RNAs induce DNA methylation at the targeted locus. c Representative genome browser view showing the DNA methylation established at Target 1 (left panel) and Target 2 (right panel) in control lines (carrying an unrelated transgene) and IR-transgenic lines. d ChIP-qPCR analysis of H3K27me3 levels in control and IR-transgenic lines at Target 1 (IR1, left panel) or Target 2 (IR2, right panel). Bar plots show the mean H3K27me3 levels from three independent lines for IR1 and 7 lines for IR2; each dot represents one independent transgenic line. A biological replicate of this experiment is shown in Supplementary Fig.5. No statistical test was performed. e ChIP-qPCR analysis of H3K27me3 levels in DRM2^WT^ or DRM2^CAT^ T1 plants transformed with IR-control, IR1 (left panel), IR2 (middle panel) or *COPIA21* (right panel) constructs. Data represent the mean of biological replicates (IR1, n=2; IR2, n=3; *COPIA21*, n=2). Each replicate corresponds to an independent pool of T1 plants. Data were normalized to the positive control *FLC*; *UBQ* serves as a negative control. No statistical test was performed. Source data are provided as a Source Data file.

To uncouple the role of DNA methylation from the role of DRM2 protein per se in this process, we introduced the hairpin construct into published lines^46^ expressing either the wild-type DRM2 (DRM2^WT^) or a catalytically inactive DRM2 (DRM2^CAT^) which localization is maintained but which cannot catalyze DNA methylation^46^. In DRM2^WT^ lines, H3K27me3 decreased as expected, whereas in DRM2^CAT^ lines, H3K27me3 levels remained comparable to the control lines (Fig. 4e, left), showing that when small RNA-mediated DNA methylation cannot be established, there is no impact on H3K27me3 levels at the target TE. Thus, it is DNA methylation, rather than DRM2 itself, that antagonizes H3K27me3 deposition.

Likewise, to verify that the gain of H3K27me3 at the DNA hypermethylated regions is the direct consequence of ectopic DNA methylation, we constructed two other inverted repeat transgenes, this time targeting two loci that gain H3K27me3 in *rdd* (*AT5TE81105* and *AT3TE23150*, i.e. LACK Target 2 and Target 3 respectively) (Fig. 4a middle and right panels, Supplementary Fig. 5a). In the RNAi lines, we observed a gain of DNA methylation in all three sequences contexts (Fig. 4c, right panel) and this time an increase in H3K27me3 at the endogenous TEs (“Target 2” and “Target 3”) targeted by RNAi and DNA methylation compared to the transgenic lines containing an unrelated transgene (Fig. 4d and Supplementary Fig. 5b right panel, Supplementary Fig. 5c). This indicates that targeting DNA methylation to a “LACK” type-TE can increase directly the recruitment of PRC2. This finding is in accordance with the striking observation that H3K27me3 deposition at the newly inserted *COPIA21* TE is dependent on DRM2 (Fig. 1c) and provides further evidence that DNA methylation can favor H3K27me3 deposition in a locus-specific manner.

Importantly, in DRM2^WT^ lines, upon inverted-transgene introduction, this gain of H3K27me3 was reproduced (Fig. 4e, right; Supplementary Fig. 5d), but it was abolished in DRM2^CAT^ lines showing that the gain of H3K27me3 is directly caused by DNA methylation rather than the presence of DRM2 itself. Accordingly, when using a similar experimental set-up as in Figure 1, a neo-inserted *COPIA21* similarly showed loss of H3K27me3 *de novo* recruitment in DRM2^CAT^ lines (Fig. 4f), confirming that it is DNA methylation and not DRM2 itself that promotes H3K27me3 at LACK.

Together, these findings demonstrate that DNA methylation can both antagonize and promote H3K27me3 deposition in a locus-specific manner.

## DISCUSSION

We previously showed that H3K27me3 recruitment and patterning at TEs share commonalities with PcG target genes, such as a partial dependency on PRC1, H2AZ variant incorporation, and the activity of JMJ histone demethylases; moreover, sequence recognition motifs such as PREs^47^ may also be involved in H3K27me3 deposition since H3K27me3 appears to be instructed by the TE sequence itself^37^. Here, by studying the establishment of H3K27me3 newly inserted TE sequences which model TE neo-insertions, we reveal a previously unappreciated dependence of PRC2 activity on DNA methylation at TEs. Specifically, while *de novo* deposition of H3K27me3 was enhanced at a neo-inserted *COPIA27* sequence, it was strikingly impaired in the *drm2* mutant at a neo-inserted *COPIA21*. As illustrated in Fig. 5a (panels 1-2), these results define two distinct modes of PRC2 recruitment at neo-inserted TEs: one in which PRC2 recruitment is promoted by low levels of DNA methylation (panel 1), and another in which PRC2 recruitment is antagonized by high DNA methylation levels at presumably PRE-rich sequences (panel 2). Together, these observations reveal a dual and context-dependent role of DNA methylation in PRC2 recruitment during silencing establishment.

**Figure 5.**
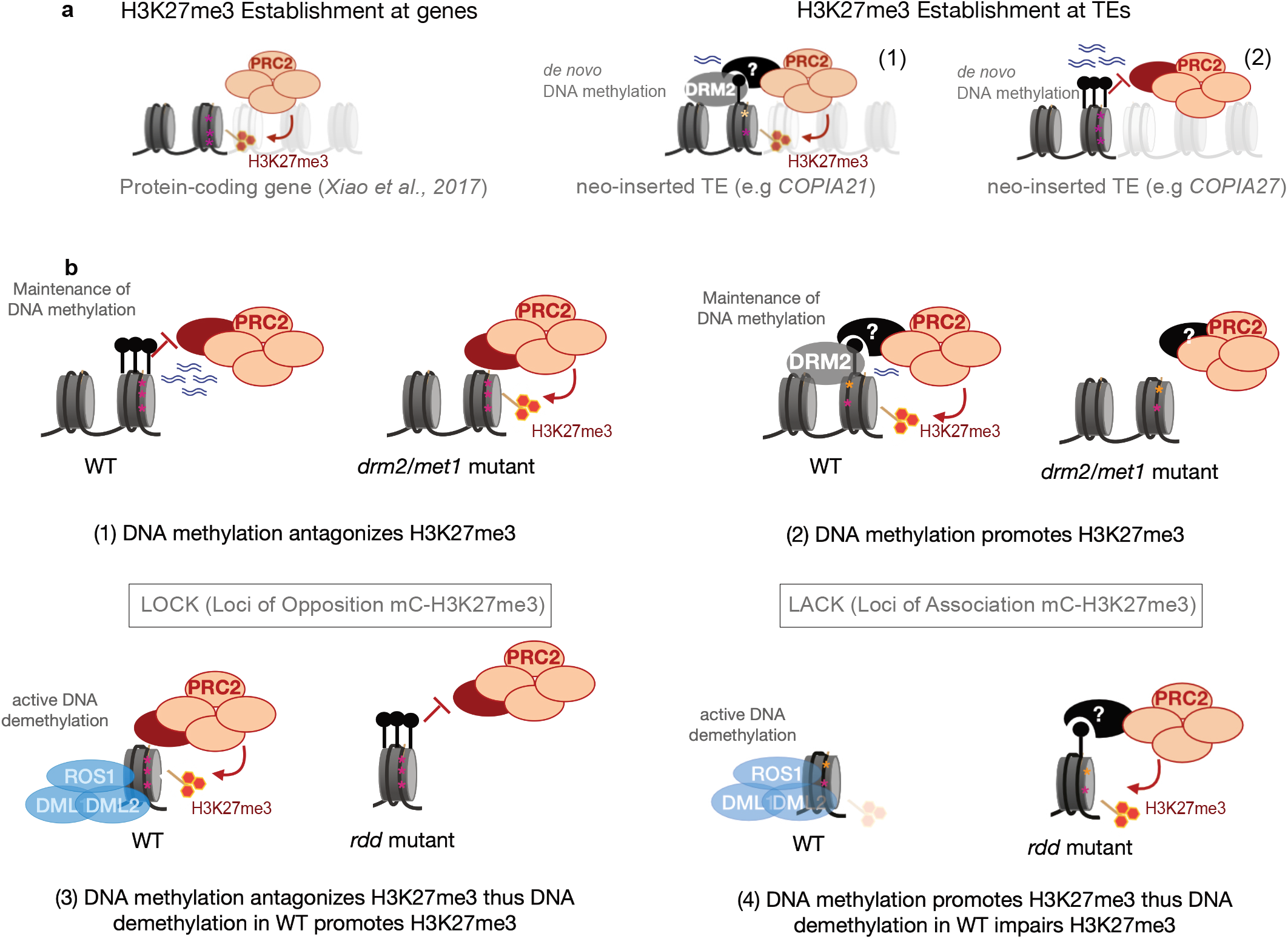
Scenario for differential epigenetic targeting of TEs, in a locus specific manner. **a** Proposed model for establishment of H3K27me3 at neo-inserted TEs. Left: PRC2 recruitment on unmethylated protein-coding genes (previously shown to be dependent on specific transcription factors (TF), Xiao et al., 2017). Purple stars: known PREs, red lollipops: H3K27me3. Right: (1) PRC2 recruitment on a neo-inserted TE sequence, lowly methylated, is DRM2-dependent. Possibly the same TF are involved or different TF needing DNA methylation/ DRM2 as a docking site to enhance their recruitment; alternatively unknown PRC2 accessory proteins with affinity to DNA methylation/DRM2 could be involved. Yellow stars: possible sequence motifs different from known canonical PREs. Right: (2) PRC2 recruitment on a neo-inserted TE sequence, highly methylated and PRE-rich, is antagonized by DNA methylation **b** Model for locus-specific rules of H3K27me3 maintenance at endogenous TEs. (1) These TEs are highly DNA methylated in WT and gain H3K27me3 marks in DNA methylation mutants (LOCK). DNA methylation presumably prevents the binding of a TF or PRC2 co-factor (red circle). Depending on the sequence motif recognized by the TF, the context of DNA methylation preventing TF binding may be different. (2) These TEs are both DNA methylated and marked by H3K27me3 in WT (LACK). DNA methylation would be needed to promote PRC2 recruitment via a TF or PRC2 co-factor (black circle) which is affine to DNA methylation and requires it as a docking site. (3) These TEs are marked by H3K27me3 only in WT, and loose H3K27me3 upon gain of DNA methylation (LOCK) in *rdd* mutant (LOCK). PRC2 recruitment is sensitive to DNA methylation, and active demethylation by DML enzymes promotes H3K27me3 recruitment at these TEs. (4) These TEs are weakly marked by H3K27me3 and DNA methylation in WT and gain H3K27me3 upon gain of DNA methylation (LACK) in *rdd*. PRC2 recruitment would be dependent on DNA methylation (as in (2)). Together, these TE-specific behaviors are associated with genetic and epigenetic signatures for PRC2 recruitment and PRC2 antagonism respectively.

Moreover, these effects are not restricted to neo-inserted sequences: at endogenous TEs we uncovered a similar dual impact of DNA methylation on PRC2 recruitment which either require DNA methylation or is antagonized by its presence (Fig. 5b) and we demonstrate that these effects are direct. We have characterized and named these two classes of loci “LACK” (“Loci of Association between Cytosine methylation and H3K27me3”) and “LOCK” (“Loci of Opposition between Cytosine methylation and H3K27me3”). As illustrated in Fig.5b, LOCK loci correspond to highly methylated TEs in which DNA methylation prevents PRC2 recruitment (panels 1 and 3). The antagonistic effect of CG methylation on H3K27me3 deposition was previously shown^31^; however, our results provide two novel insights into that aspect. First, DRM2-mediated non-CG methylation can also antagonize PRC2 recruitment (panel 1), which is consistent with recent H3K27me3 profiles of rice and maize RdDM mutants^50,51^. Second, we demonstrate a novel role for DNA demethylases in promoting PRC2 recruitment (panel 3). This result is in line with previous observations that ROS1 targets are enriched in H3K27me3^43^ and could provide an explanation as to why mammalian TET enzymes are enriched at the hypomethylated DNA promoters of PcG targets^52^. By contrast, LACK loci are both DNA methylated and marked by H3K27me3, and require DNA methylation, in all contexts and at low levels, to promote H3K27me3 deposition (panels 2 and 4). Interestingly, this relationship was also recently reported in rice genes where H3K27me3 and non-CG methylation can colocalize^48,49^ although levels of DNA methylation were not examined in this case.

Thus, to reconcile the apparently opposite effects of DNA methylation on PRC2 recruitment, our data support a locus-specific model for H3K27me3 maintenance at endogenous TEs. Our analyses rule out that loss of H3K27me3 upon loss of DNA methylation is due to transcriptional reactivation. Instead, they suggest a mode of PRC2 recruitment determined by a combination of specific genetic and epigenetic signatures specific to each target.

First, our analyses indicate that a certain threshold of DNA methylation could contribute to determine whether H3K27me3 deposition is antagonized (high DNA methylation) or promoted (low DNA methylation). This is also consistent with neo-inserted *COPIA21* displaying less DNA methylation than neo-inserted *COPIA27*. Second, we found that LACK are associated with different chromatin states than LOCK, the pertinence of which should be addressed in future endeavours. For instance, at some loci, H2Aub could be involved in the recruitment of the DNA methylation sensitive K27me3 demethylase REF6 which can modulate H3K27me3 levels in a DNA methylation-dependent manner^53,54^, but this remains to be tested. Active DNA demethylation can further modulate these states as illustrated by Fig. 5b (panels 3-4): at LOCK, DNA demethylation promotes H3K27me3 deposition (panel 3), whereas at LACK, DNA demethylation impairs PRC2 recruitment (panel 4). This highlights the dynamic interplay between DNA methylation turnover and PRC2 targeting and could further suggest that some LOCK and LACK may contribute to the regulation of nearby gene. This would be consistent with the proposed function of ROS1 in enabling gene responsiveness through DNA methylation removal at cis-regulatory TEs ^22,57^; this possibility warrants future investigation.

In addition, presumably genetic signatures contribute to differential PRC2 recruitment. Notably, we found that LOCK were significantly enriched in several previously described PREs compared to LACK: this could provide an explanation as to why PRC2 needs additional features, such as DNA methylation at low levels, to be recruited efficiently to LACK . This also points to the existence of a set of transcription factor (TF) or PRC2 co-factors that may recognize specific sequence patterns, that display different affinities for DNA methylation and that need to be identified in the future.

This scenario does not exclude physical interactions between PRC2 and DRM2 or the DNA methylation machineries in a locus-specific manner or in a context-specific manner (for example, during neo-insertion events). However, our results using a *drm2* catalytic mutant demonstrate that it is DNA methylation, not DRM2 protein itself, that influences PRC2 recruitment. Moreover, Alphafold predictions indicate that the direct interactions between DRM2 and the various PRC2 subunits are unlikely (Supplementary Fig. 6, Supplementary Data 1). Therefore, we propose that any potential physical association between these machineries is likely to be indirect, mediated through DNA methylation recognition, and involving bridging factors—such as transcription factors recognizing PREs or methyl-DNA–binding proteins—that remain to be identified (Fig. 5b). Given their role in PRC2 recruitment^28^, it is also possible that non-coding RNAs associated with DNA methylation could also be important in the positive crosstalk between DNA methylation and PRC2. In that respect, the transgenic system that we have established provides a framework to dissect the complex interactions between H3K27me3 and DNA methylation during establishment and maintenance throughout generations at a single TE copy in future work.

The interconnection that we revealed between *de novo* DNA methylation and deposition of H3K27me3 on newly inserted TEs argues for active cooperation between the two marks at this stage of the TE life cycle. This raises the question as to why such cooperation would take place when a novel copy has just integrated into the genome. When they are just inserted into the genome, retrotransposons are DNA hypomethylated. The targeting of PcG at this stage could thus allow rapid silencing of the element while DNA methylation is being established progressively throughout successive generations, especially if there is a low amount of homologous small RNA that can establish *de novo* DNA methylation. One exciting question to address in the future is whether H3K27me3 persists after one round of reproduction or if DNA methylation quickly becomes dense enough to antagonize H3K27me3 in subsequent generations. If this is the case, H3K27me3 could just be a transient silencing form established quickly by the plant to compensate for low *de novo* DNA methylation in the first generation(s) after TE neo-insertion. Our study thus points to a synergy between the two silencing pathways, a concept that has also recently emerged at other types of repeats^55,56^.

By connecting PRC2 to DNA methylation at TEs, our work in Arabidopsis provides important insights into the separation and specialization of the two major silencing pathways throughout eukaryotic evolution. We previously proposed that PcG was an ancestral system of TE silencing based on H3K27me3 being the dominant mark at TEs in unicellular organisms or ancestral plants. Furthermore, in ciliates, a small RNA-guided Enhancer of Zeste (Ezl-1, the conserved catalytic subunit of PRC2) deposits both H3K27me3 and H3K9 methylation^23,58,59^, which either reflects ciliate-specific catalytic activity or suggests that ancestral PRC2 has both activities. We propose that with the evolution of multicellularity and the need for a dynamic system to control developmental transitions, the PcG and H3K9me2/DNA methylation pathways may have specialized for the silencing of genes and TEs, respectively. Our present results nevertheless show that the major silencing pathways in eukaryotes maintain mechanistic connections despite specialization for different functions in higher plants, which is likely to help their functional cooperation in specific contexts. Such interconnections may exist in other kingdoms, as suggested by the small RNA-driven deposition of H3K27me3 in *C. elegans*^60,61^ or the existence of AEBP2, a mammalian PRC2 cofactor that requires DNA methylation for its activity in vitro^62^. Conversely, DNA methylation could be dependent on PRC2, as previously suggested^63,64^, and it would be interesting to investigate this possibility in plants. Future work needs to further decipher the connections between PcG and H3K9/DNA methylation and how they evolved from unicellular to multicellular organisms. This should undoubtedly shed light on the evolution of silencing pathways in eukaryotes and how they shape host genome regulation.

## METHODS

### Plant material and growth conditions

All the experiments were conducted on *A. thaliana* on ½ MS plates under short light-day conditions (8-h light/16-h dark photoperiod at 22°C). For the transgenic plants shown in Fig. 1, 4- week-old rosette leaves were pooled and collected for further ChIP-seq and BS-seq analysis.

### Mutant lines

We used the *drm1-2 drm2-2* double mutant (Salk-031705, Salk-150863)^5^ , the *met-1* mutant (NASC <u>N16394</u>)^65^ and the *ros1 dml2 dml3* triple mutant (Col-0 background, derived from Salk_045303, Salk_131712, Salk_056440, respectively)^13^

### Generation of transgenic lines

The *COPIA21* and *COPIA27* TE sequence were synthesized and cloned and inserted into pUC57 via GenScript. *COPIA21* and *COPIA27* sequences were subsequently cloned and inserted into pCAMBIA3300. The plants were subsequently transformed via *Agrobacterium tumefaciens* floral dip^66^. For the experiments shown in Fig. 1, the transgenic plants were selected on Basta, transferred in soil after 2 weeks and 4 weeks old rosettes leaves were pooled from 18-20 T1 plants to dilute position effects, to perform ChIP-seq and BS-seq (on the same ground tissue).

RNAi lines were obtained by cloning approximately 250 bp fragments in an inverted orientation via the pFRN vector^74^. The plants were subsequently transformed via *Agrobacterium tumefaciens* floral dip^66^, and T1 plants were selected *in vitro* on Kanamycin.

### Chromatin immunoprecipitation (ChIP) and ChIP‒qPCR/sequencing analyses

ChIP experiments were conducted in WT or appropriate mutant lines via an anti-H3K27me3 antibody (Millipore 07-749 for Figure 1,Diagenode C15410069 for the remainder of the paper, 4ug/IP). IP and INPUT DNA were eluted, purified and sequenced (100 bp paired-end; Illumina) by BGI. Reads were mapped via BWA^67^ onto TAIR10 *A. thaliana.* Genomic regions significantly marked by H3K27me3 were identified via MACS2^68^ and genes or TEs overlapping these regions were obtained via bedtools^69^. Heatmaps and plotprofiling were generated via bedtools computeMatrix to create a score matrix, and plotHeatmap was used to generate a graphical output of the matrix. For ChIP-seq analyses in *rdd* mutant, regions inherited from Ws-2 *ros1* and *dml2* mutants (Ws background) after backcrossing into Col-0 were excluded from analysis as previously described^14^. Primers used are listed in Supplementary Table 1.

### Bisulfite-sequencing analyses

DNA was extracted using a standard cetyl trimethylammonium bromide (CTAB)-based protocol. Adapter and low-quality sequences were trimmed via Trimming Galore 0.6.5. Mapping was performed on the TAIR10 genome annotation via Bismark v0.22.2^70^ and Bowtie2^71^.

DNA methylation at *COPIA21* transgenic copies was estimated using the number of transgenic copies (based on read counts in transgenic lines compared to the non-transgenic lines), the total number of *COPIA21* copies (endogene + transgenes), the DNA methylation level of endogenous *COPIA21* in non-transgenic lines and the DNA methylation level of *COPIA21* in transgenic lines. We used the following formula, where *x* stands for the DNA methylation level in a given cytosine context at a single transgenic copy, and then presented on the y axis in %:

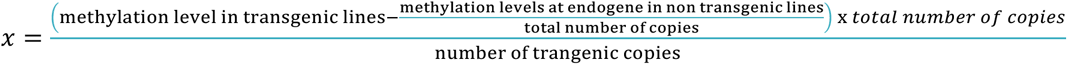

The same approach was used for *COPIA27*

### Analyzed data sets

The public datasets were downloaded from NCBI GEO or ENA using the specified accession numbers: H3K27me3 ChIP-seq in wild-type and *COPIA21* transgenic lines (PRJEB83093)^37^. H3K27me3 ChIP-seq in *ddm1* (<u>PRJEB34363</u>)^32^. Small-RNAs sequencing in wild-type (GSE224761)^38^. BS-seq Col-0, *drm2 and met1* (<u>GSE39901</u>)^72^, and BS-seq Col-0 and *rdd* (GSE33071)^73^. ChIP-seq of WT histone marks, methylation (<u>GSE226469</u>)^39^, and acetylation (GSE79524)^40^.

### Statistics and reproducibility

The relevant information is described in the figures legends. No statistical method was used to predetermine sample size. No data were excluded from the analyses. The experiments were not randomized. The investigators were not blinded to allocation during experiments and outcome assessment.

## DATA AVAILABILITY

All sequencing datasets generated in this study are publicly available in the Sequence Read Archive (SRA) repository under the accession number <u>PRJNA1464127</u>. The processed files associated to raw data are available on Gene Expression Omnibus (GEO) under the accession number GSE331299. Source data are provided with this paper. Lists of LACK and LOCK defined in *drm2*, *rdd*, or *met1* are in Supplementary Data 2.

## Acknowledgements

This work was funded by French State Grant ANRJCJC (ANR-19-CE12-0033-01 to A.D.) and has benefited from a grant Saclay-Plant Sciences (SPS) (ANR-17-EUR-0007, EUR SPS-GSR) under a France 2030 program (ANR-11-IDEX-0003) and SEVE Paris-Saclay Doctoral School (PhD Fellowship to M.K.E.). We thank the Genome Biology Department of Institut de Biologie Intégrative de la Cellule (I2BC) for support and discussions. We thank the services and platforms of the Institut de Biologie Intégrative de la Cellule (I2BC) for excellent technical support and in particular Véronique Couvreux for excellent plant care. We thank N. Bouché for his help with some bioinformatic analyses and Stéphane Pelletier and Mathieu Rougemaille for technical help during the revisions. We thank our colleagues of the CNRS Groupements de Recherche “Epiplant” (GDR2027) and “Mobil’ET” (GDR3546) for discussions.

## Author information

### Author notes

These authors contributed equally: Valentin Hure and Maghnia Kawter Elmir

### Contributions

V.H. and A.D. designed the study. V.H and M.K.E performed most of the experiments; V.H. performed computational analyses. F.P.P performed ChIP-seq analyses et DNA extraction for Figure 1. R.G performed AlphaFold predictions for Supplementary Figure 6 as part of the revisions. V.H. and A.D. prepared the manuscript. V.H., M.KE, and A.D revised the manuscript.

**Corresponding author:** Correspondence to Angélique Déléris

Competing interests:

The authors declare no competing interests

---

**Supplementary Figure 1.**
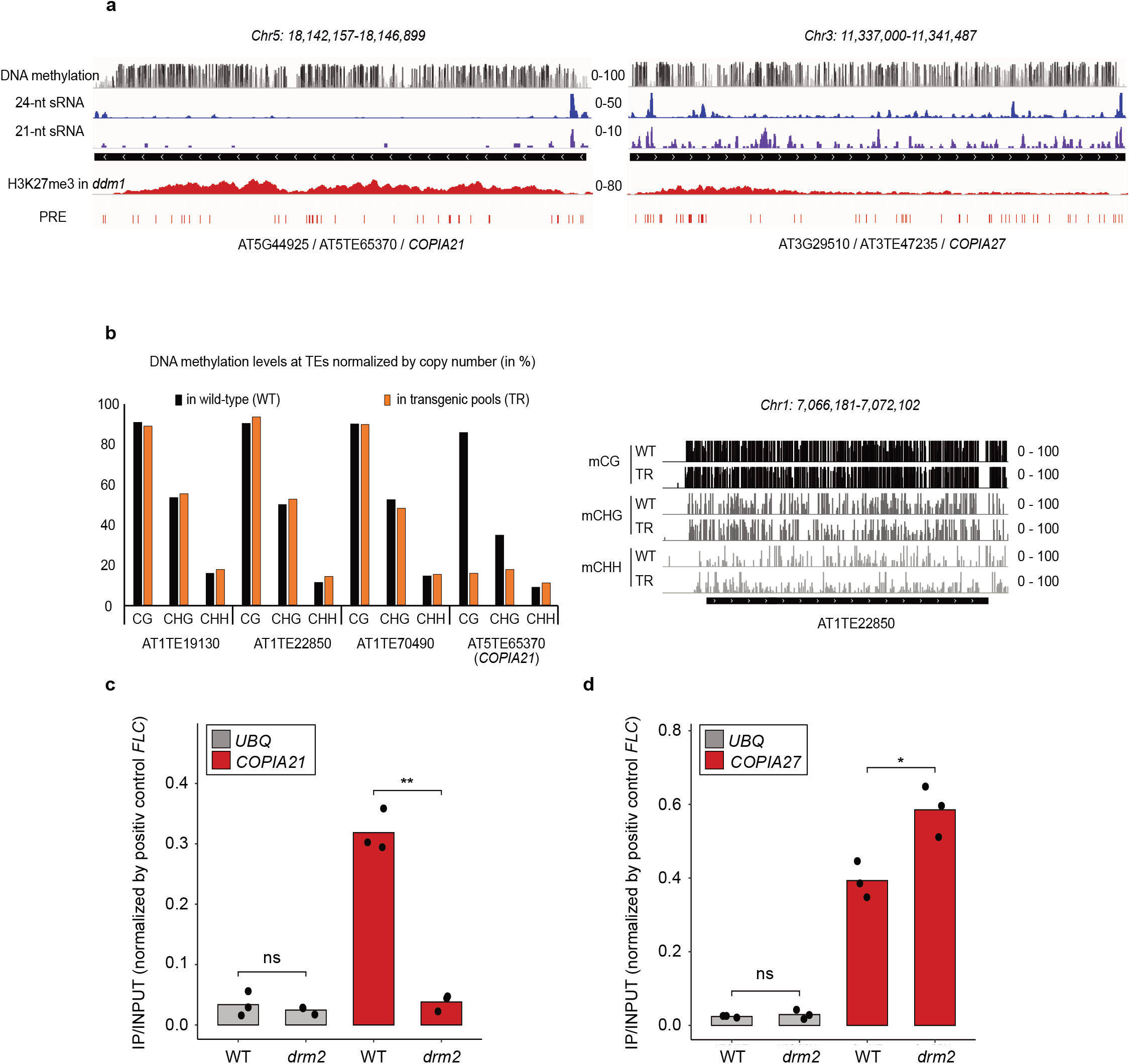
Role of DRM2 in the establishment of *de novo* H3K27me3 at a neo-inserted TE transgenic sequence. **(a)** Representative genome browser view of DNA methylation (CG: black, CHG: dark grey, CHH: light grey), 24-nt small RNAs levels (blue) and 21/22-nt small RNAs levels (purple) in wild-type, H3K27me3 levels in ddm1 mutant and Polycomb-Responsive Elements (PREs) (red) at *COPIA21* (**left panel**) and *COPIA27* (**right panel**). **(b) Left panel**: Bar plot showing the DNA methylation levels in each cytosine context at different endogenous TEs in WT (black bars) and in transgenic pools (orange bars). **Right panel**: Representative genome browser view of DNA methylation status (CG: black, CHG: dark grey, CHH: light grey) at *AT1TE22850* in wild-type (WT) and in transgenic pools (TR). **(c)** Bar plot showing H3K27me3 levels assessed by ChIP-qPCR at *COPIA21* in wild-type and *drm2 COPIA21* transgenic lines. IPs were normalized by the input and then normalized by the positive control *FLC*. Three independent biological replicates are shown. **(d)** Bar plot showing H3K27me3 levels assessed by ChIP-qPCR at *COPIA27* in wild-type and *drm2 COPIA27* transgenic lines. IPs were normalized by the input and then normalized by the positive control *FLC*. Three independent biological replicates are shown.

**Supplementary Figure 2.**
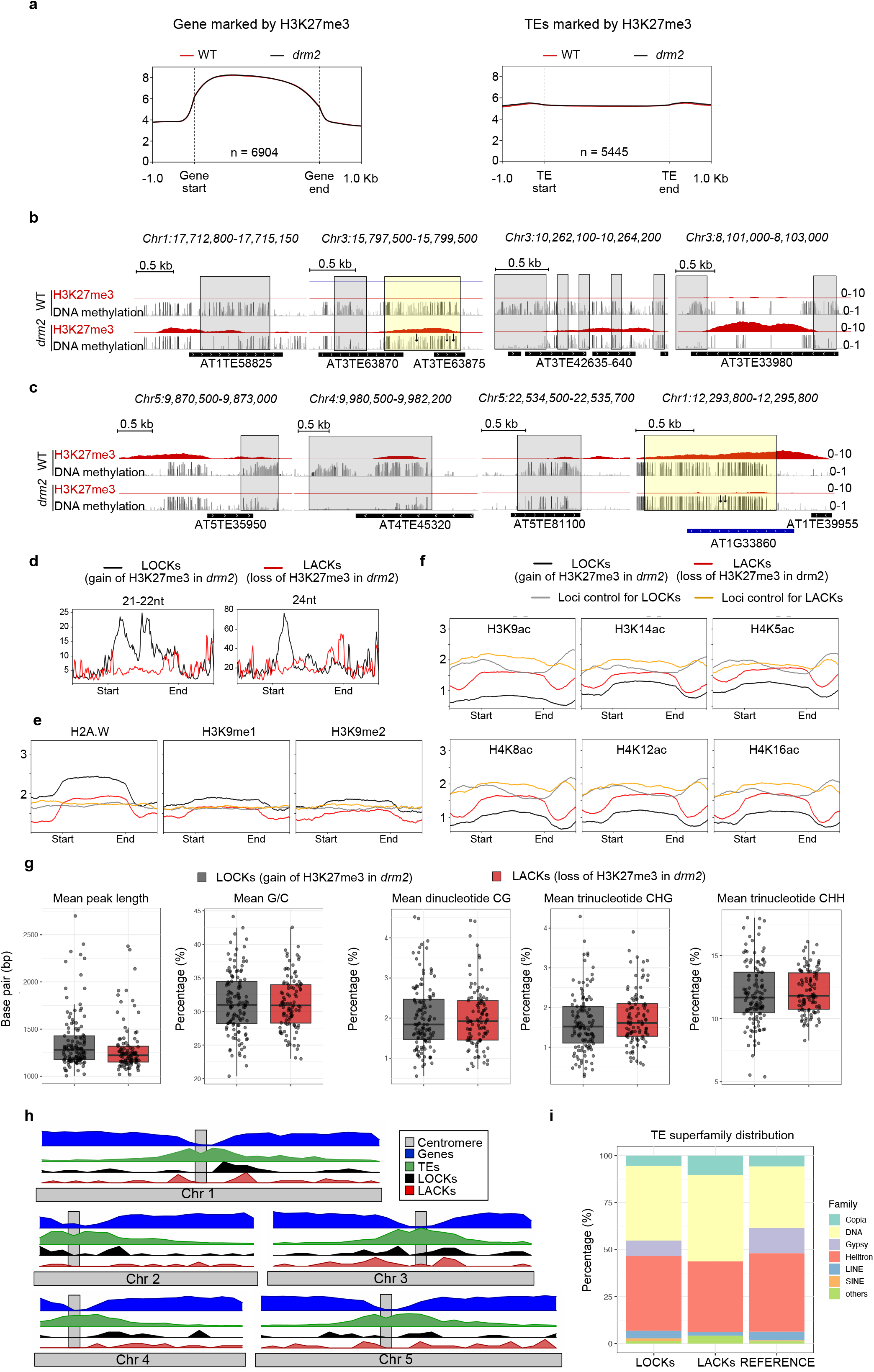
Role of DRM2 in the recruitment of H3K27me3 at endogenous loci. **(a)** Metagenes showing H3K27me3 level for all genes (**left panel**) and all TEs (**right panel**) marked by H3K27me3, in WT and *drm2* mutant. **(b)** Representative genome browser views showing H3K27me3 ChIP-seq and DNA methylation BS-seq data in WT and *drm2* mutant. Gain of H3K27me3 in *drm2* compared to WT is either associated with previously published DMRs (grey boxes) (*Stroud et al, 2013*) or DNA methylated regions that lose several methylated cytosines (yellow box; the arrows indicate the loss of several mCHH). **(c)** Representative genome browser views showing H3K27me3 ChIP-seq and DNA methylation BS-seq data in WT and *drm2* mutant. Loss of H3K27me3 in *drm2* compared to WT is either associated with previously published DMRs (grey boxes) (*Stroud et al, 2013*) or DNA methylated regions that lose several methylated cytosines (yellow box; the arrows indicate the loss of several mCHH). **(d)** Metaplots showing 21/22-nt and 24-nt small RNAs profiles at LOCKs (loci that gain H3K27me3 in *drm2*, black) and at LACKs (loci that loose H3K27me3 in *drm2*, red). **(e)** Metaplots showing H2A.W variant, H3K9me1 and H3K9me2 profiles at LOCKs (black), at LACKs (red) and at associated control regions (random loci of the same length in the genome). **(f)** Metaplots showing H3K9acetylation, H3K14ac, H4K5ac, H4K8ac, H4K12ac and H4K16ac profiles at LOCKs (black), at LACKs (red) and at associated control regions (random loci of the same size in the genome). **(g)** Box plots showing the mean of peak length, the mean of C and G nucleotides, the mean of CG dinucleotides, the mean of CHG trinucleotides and the mean of CHH trinucleotides at LOCKs (black) and LACKs (red). **(h)** Genomic distribution of the LOCKs (black), the LACKs (red), the genes (blue), the TEs (green) and the centromeric region (grey). **(i)** Stacked bar plot showing the distribution of TE superfamilies in LOCKs (black), LACKs (red), and in the whole genome as ‘REFERENCE’.

**Supplementary Figure 3.**
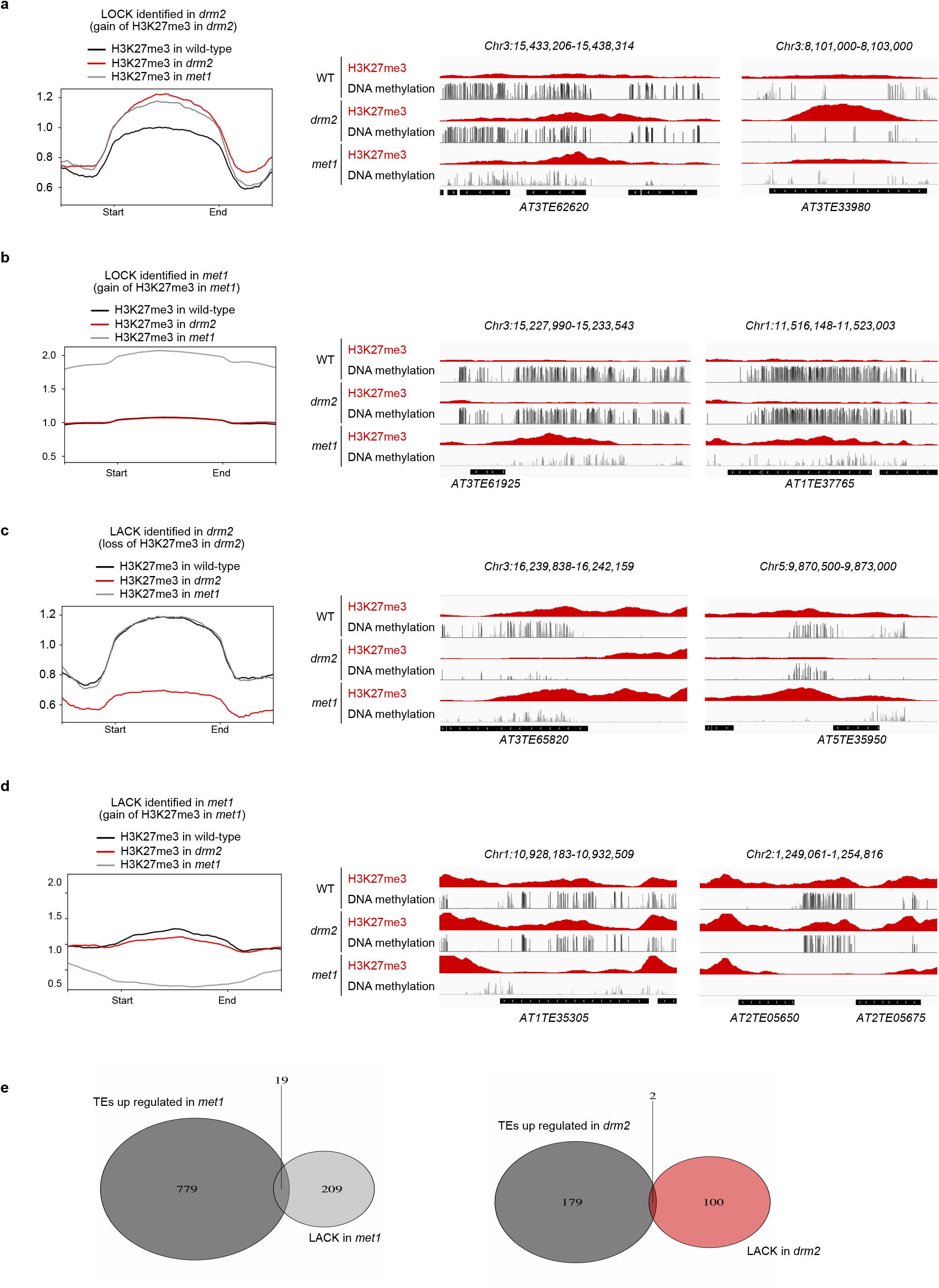
Role of MET1 in the recruitment of H3K27me3 at endogenous loci. loci. **(a)** Metaplots showing H3K27me3 profiles in wild-type (black), *drm2* (red), and *met1* (grey) at LOCKs identified in *drm2* (loci that gain H3K27me3 in *drm2*). Representative genome browser views of LOCKs identified in *drm2*, showing H3K27me3 ChIP-seq and DNA methylation BS-seq data in WT, *drm2* and *met1* mutant. **(b)** Metaplots showing H3K27me3 profiles in wild-type (black), *drm2* (red), and *met1* (grey) at LOCKs identified in *met1* (loci that gain H3K27me3 in *met1*). Representative genome browser views of LOCKs identified in *met1*, showing H3K27me3 ChIP-seq and DNA methylation BS-seq data in WT, *drm2* and *met1* mutant. **(c)** Metaplots showing H3K27me3 profiles in wild-type (black), *drm2* (red), and *met1* (grey) at LACKs identified in *drm2* (loci that loose H3K27me3 in *drm2*). Representative genome browser views of LACKs identified in *drm2*, showing H3K27me3 ChIP-seq and DNA methylation BS-seq data in WT, *drm2* and *met1* mutant. **(d)** Metaplots showing H3K27me3 profiles in wild-type (black), *drm2* (red), and *met1* (grey) at LACKs identified in *met1* (loci that loose H3K27me3 in *met1*). Representative genome browser views of LACKs identified in *met1*, showing H3K27me3 ChIP-seq and DNA methylation BS-seq data in WT, *drm2* and *met1* mutant. **(e)** Left panel: Venn diagram showing intersection between up-regulated TEs in *met1* (*Zhao et al, 2022*) and our LACKs identified in *met1*. Right panel: Venn diagram showing intersection between up-regulated TEs in *drm2* (*Stroud et al, 2014*) and our LACKs identified in *drm2*.

**Supplementary Figure 4.**
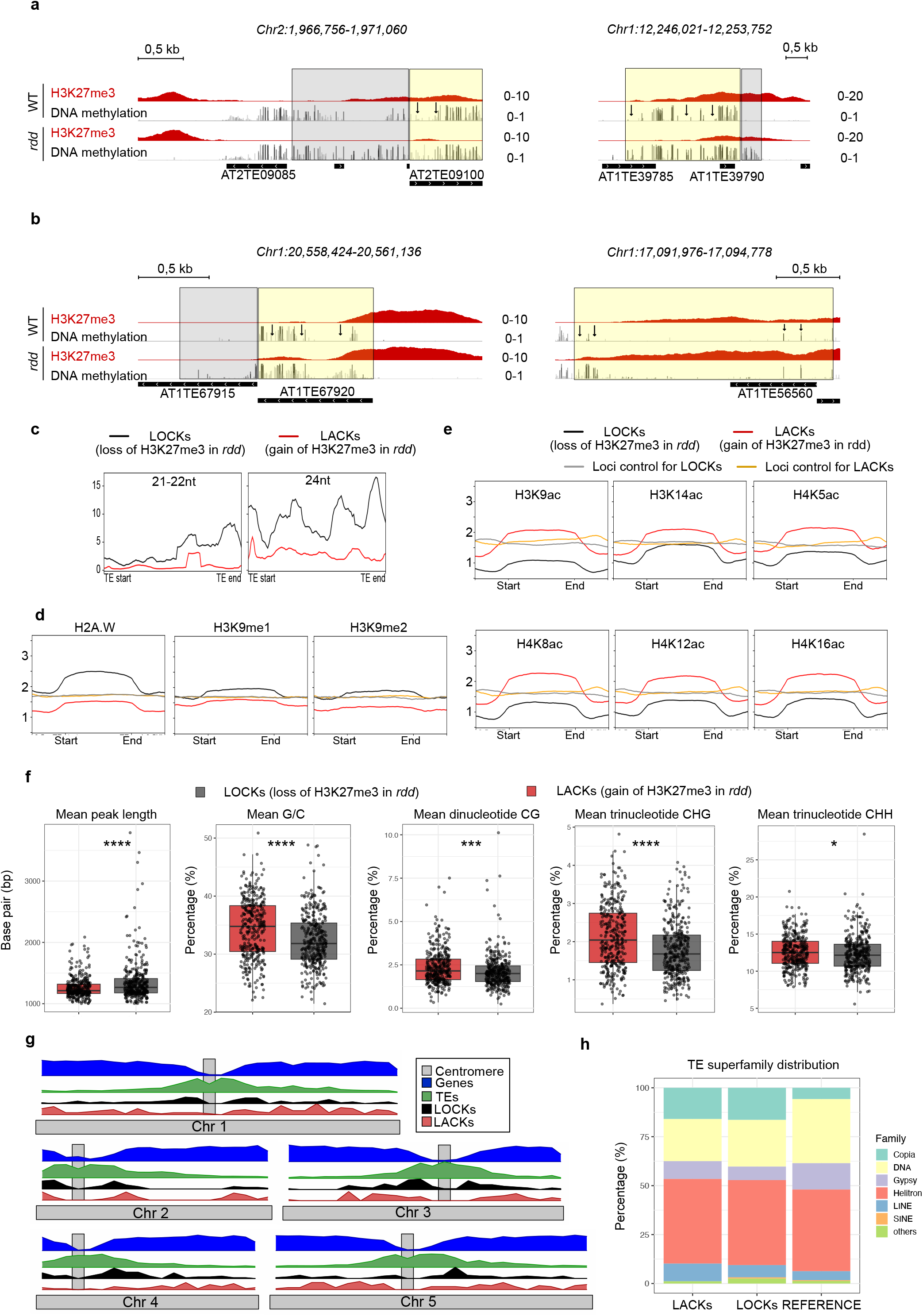
DNA demethylation impacts H3K27me3 patterning at endogenous loci. **(a)** Representative genome browser views showing H3K27me3 ChIP-seq and DNA methylation BS-seq data in WT and *rdd* mutant. Loss of H3K27me3 in *rdd* compared to WT is either associated with previously published DMRs (grey boxes) (*Duan et al, 2015*) or DNA methylated regions that gain several methylated cytosines (yellow box; the arrows indicate the gain of several mCHH). **(b)** Representative genome browser views showing H3K27me3 ChIP-seq and DNA methylation BS-seq data in WT and *rdd* mutant. Gain of H3K27me3 in *rdd* compared to WT is either associated with previously published DMRs (grey boxes) (*Duan et al, 2015*) or DNA methylated regions that gain several methylated cytosines (yellow box; the arrows indicate the gain of several mCHH). **(c)** Metaplots showing 21/22-nt and 24-nt small RNAs profiles at LOCKs (loci that lose H3K27me3 in *rdd*, black) and at LACKs (loci that gain H3K27me3 in *rdd*, red). **(d)** Metaplots showing H2A.W variant, H3K9me1 and H3K9me2 profiles at LOCKs (black), at LACKs (red) and at associated control regions (r random loci of the same length in the genome). **(e)** Metaplots showing H3K9acetylation, H3K14ac, H4K5ac, H4K8ac, H4K12ac and H4K16ac profiles at LOCKs (black), at LACKs (red) and at associated control regions (random loci of the same size in the genome). **(f)** Box plots showing the mean of peak length, the mean of C and G nucleotides, the mean of CG dinucleotides, the mean of CHG trinucleotides and the mean of CHH trinucleotides at LOCKs (black) and LACKs (red). **(g)** Genomic distribution of the LOCKs (black), the LACKs (red), the genes (blue), the TEs (green) and the centromeric region (grey). **(h)** Stacked bar plot showing distribution of TE superfamilies in LOCKs (black), LACKs (red), and in the whole genome as ‘REFERENCE’.

**Supplementary Figure 5.**
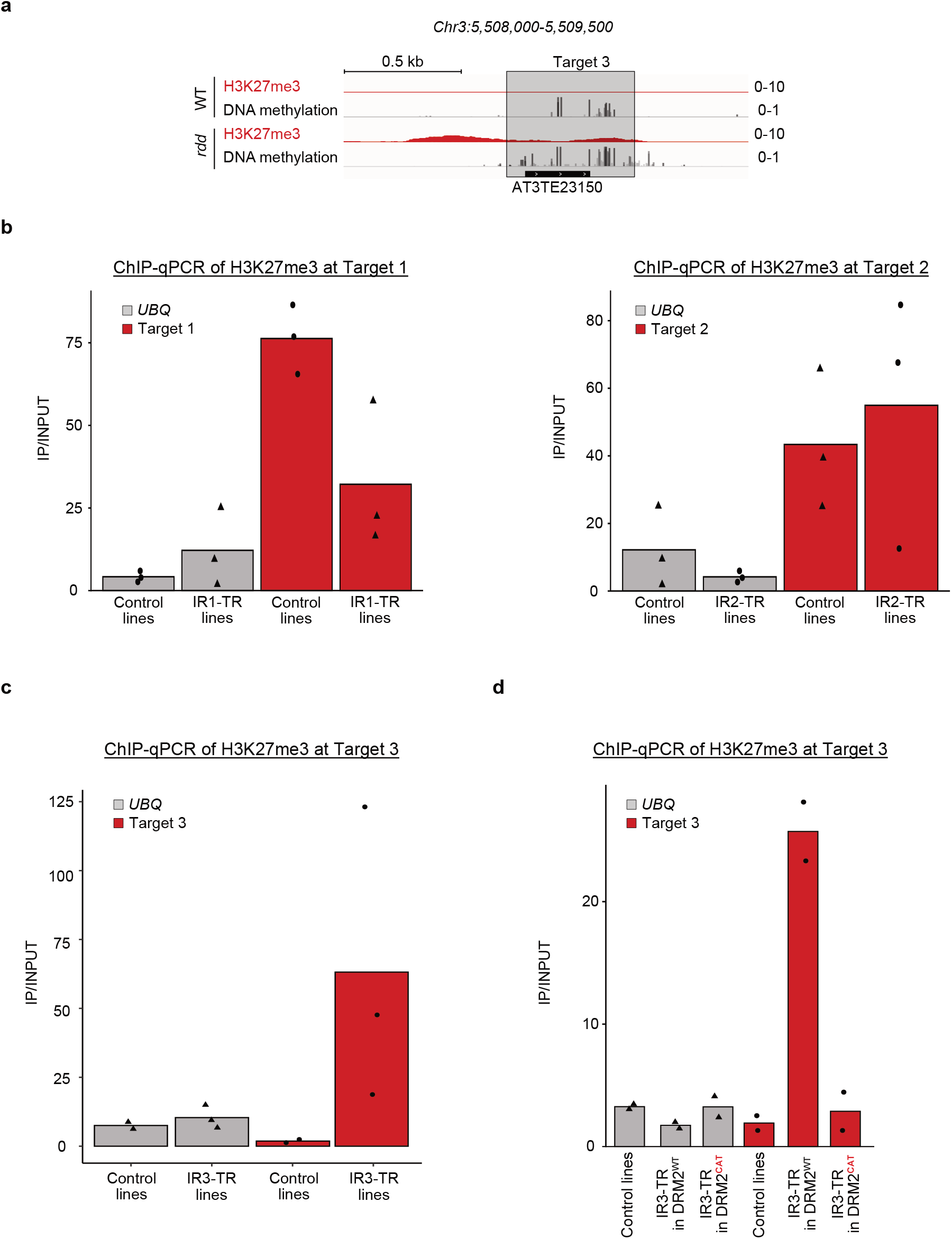
Targeted DNA methylation by Inverted-Repeats (IR) reveals a dual role of DNA methylation on H3K27me3 patterning at endogenous TEs. (a) Representative genome browser view showing H3K27me3 and DNA methylation in WT and *rdd* mutant. The region highlighted in grey corresponds to a DMR that was cloned as an inverted-repeat (IR) for the experiments shown in panels c and d. (b) ChIP-qPCR analysis of H3K27me3 levels in control and IR-transgenic lines at Target 1 (IR1, left panel) and Target 2 (IR2, right panel). This panel shows a biological replicate of the experiment presented in Fig.4 (ChIP replicate on three lines). Bar plots show the mean H3K27me3 levels from three independent control and transgenic lines. Each dot represents one transgenic line. Biological replicates are shown separately in Fig.4 and Fig.S4 due to differences in ChIP efficiencies between the two experimental batches. (c) ChIP-qPCR analysis of H3K27me3 levels in control and IR-transgenic lines at Target 3 (IR3). Bar plots show the mean H3K27me3 levels from three independent IR3 lines with each dot representing one line. (d) ChIP-qPCR analysis of H3K27me3 levels in DRM2^WT^ or DRM2^CAT^ T1 plants transformed with IR-control or IR3 constructs (pools of T1). Data were normalized to the positive control *FLC*; *UBQ* serves as a negative control.

**Supplementary Figure 6.**
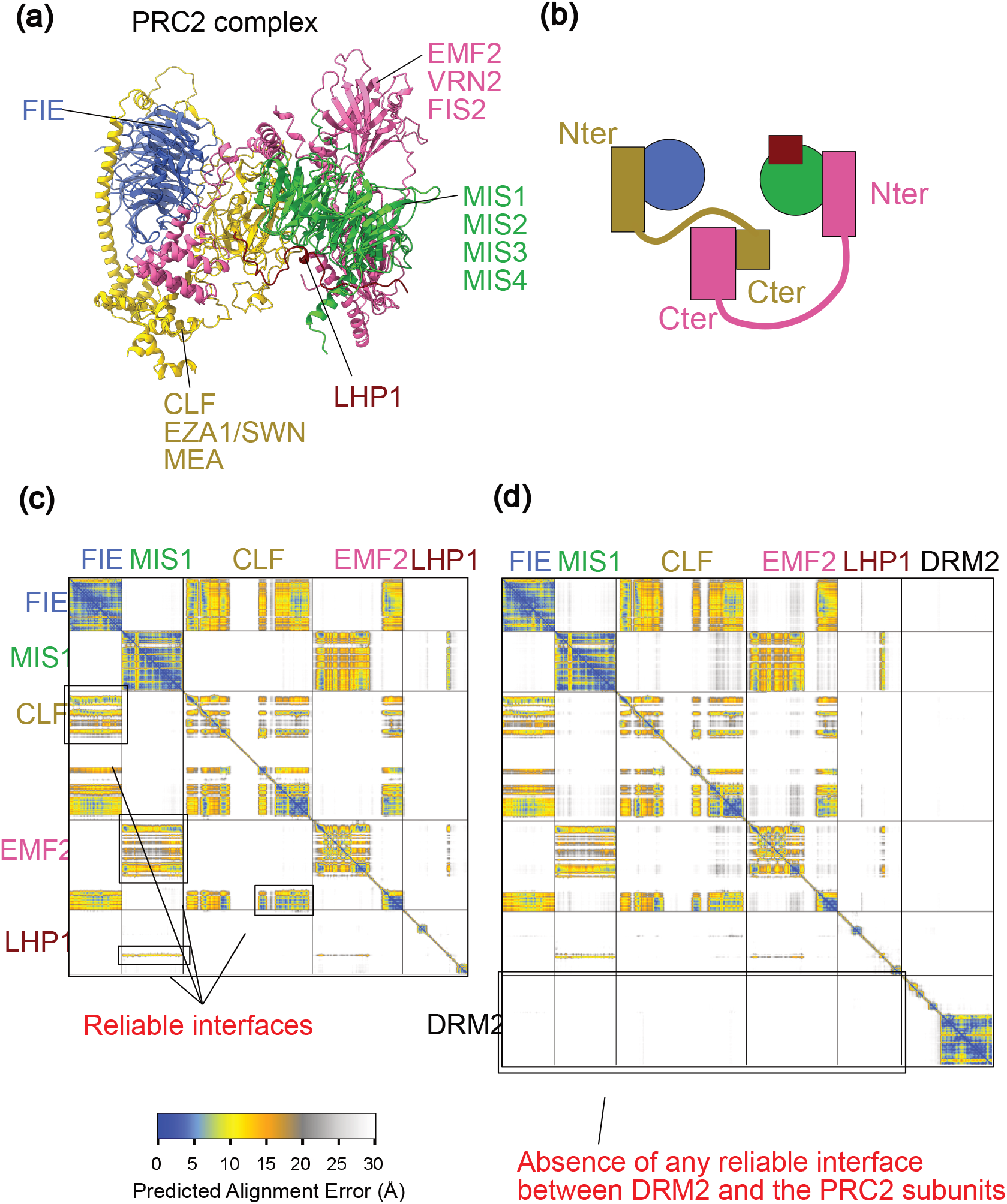
AlphaFold3 predictions reveal no reliable interfaces between DRM2 and PRC2 subunits. (**a**) Structural model of the Arabidopsis PRC2 complex, highlighting the core subunits FIE (blue), CLF (yellow) (can also be substituted by EZA1/SWN or MEA) , the MSI1 subunit (green) (can also be substituted by MSI2, MSI3 or MSI4), EMF2 (magenta) (can also be substituted by VRN2 or FIS2) and LHP1 (red). (**b**) Schematic representation of the N-terminal (Nter) and C-terminal (Cter) regions of PRC2 components, illustrating the relative positions of the interfaces and flexible regions. (**c**) Predicted Alignment Error (PAE) heatmap of the PRC2 subunits alone, showing multiple high-confidence interaction interfaces (blue/orange blocks). Arrows indicate well-supported interfaces recovered by AlphaFold3. (**d**) PAE heatmap of PRC2 subunits modelled together with DRM2, showing that DRM2 does not form any reliable interface with any PRC2 subunit (large white/grey region at the bottom right). Color scale: lower PAE values (blue) indicate high-confidence residue-residue distances, whereas higher PAE values (yellow/grey) reflect low-confidence or flexible regions.

## Notes

### Competing Interest Statement

The authors have declared no competing interest.

### Summary of Updates

First, we now provide an extensive genome-wide analysis identifying the genetic and epigenetic features that distinguish the two classes of loci at which DNA methylation either promotes H3K27me3 (now referred to as LACK for Loci of Association between Cytosine methylation and K27me3) or antagonizes H3K27me3 (now referred to these as LOCK for Loci of Opposition between Cytosine methylation and K27me3). These analyses include chromatin marks, histone variants, sequence composition and length, TE family distribution, genomic distribution, and siRNA profiles. They are further supported by experimental testing of an additional COPIA element in the neo-insertion assay. Second, we have directly tested the mechanistic basis of the relationship between DNA methylation and PRC2 recruitment: through the use of a catalytically inactive DRM2 mutant, we demonstrated that DNA methylation itself, rather than the DRM2 protein ,is required for H3K27me3 deposition. This is further supported by analyses in another DNA methyltransferase mutant. These additions strengthen the conceptual and mechanistic conclusions of the manuscript.

